# Network analysis of whole-brain fMRI dynamics: A new framework based on dynamic communicability

**DOI:** 10.1101/421883

**Authors:** Matthieu Gilson, Nikos E. Kouvaris, Gustavo Deco, Jean-François Mangin, Cyril Poupon, Sandrine Lefranc, Denis Rivière, Gorka Zamora-López

**Author notes:** Corresponding author; Universitat Pompeu Fabra, C/ Ramon de Trias Fargas, 25-27, Barcelona 08005, Spain.

## Abstract

Neuroimaging techniques such as MRI have been widely used to explore the associations between brain areas. Structural connectivity (SC) captures the anatomical pathways across the brain and functional connectivity (FC) measures the correlation between the activity of brain regions. These connectivity measures have been much studied using network theory in order to uncover the distributed organization of brain structures, in particular FC for task-specific brain communication. However, the application of network theory to study FC matrices is often “static” despite the dynamic nature of time series obtained from fMRI. The present study aims to overcome this limitation by introducing a network-oriented analysis applied to whole-brain effective connectivity (EC) useful to interpret the brain dynamics. Technically, we tune a multivariate Ornstein-Uhlenbeck (MOU) process to reproduce the statistics of the whole-brain resting-state fMRI signals, which provides estimates for MOU-EC as well as input properties (similar to local excitabilities). The network analysis is then based on the Green function (or network impulse response) that describes the interactions between nodes across time for the estimated dynamics. This model-based approach provides time-dependent graph-like descriptor, named communicability, that characterize the roles that either nodes or connections play in the propagation of activity within the network. They can be used at both global and local levels, and also enables the comparison of estimates from real data with surrogates (e.g. random network or ring lattice). In contrast to classical graph approaches to study SC or FC, our framework stresses the importance of taking the temporal aspect of fMRI signals into account. Our results show a merging of functional communities over time (in which input properties play a role), moving from segregated to global integration of the network activity. Our formalism sets a solid ground for the analysis and interpretation of fMRI data, including task-evoked activity.

## 1. Introduction

The study of the brain network has attracted much attention in recent years as a collective attempt to understand how distributed and flexible cognitive functions operate. A large body of data-driven studies has focused on the interpretation of brain connectivity measured by 5 structural and functional magnetic resonance imaging (sMRI and fMRI, respectively); for a review see [1]. A particular focus [2] has been on the relationship between the structural connectivity (SC), which is the architecture of connections between brain regions, and functional connectivity (FC), the correlation structure of the observed fMRI//BOLD activity. Initially, SC and FC were investigated using statistical descriptors designed for graphs such as the degree distribution and clustering coefficient [3, 4, 5]. Brain areas can then be characterized such as highly connected hubs that are hypothesized to centralize information distributed across the brain [6, 7]. Following, the concept of communicability was proposed as a model of interactions in a graph to link the network structure to the pairwise functional associations of nodes [8]. In essence, graph communicability takes into account indirect paths —not only shortest paths— in the definition of interactions between pairs of nodes, which is important to evaluate global effects in recurrently-connected networks [9]. It has then been used to derive measures that describe the roles for nodes in networks [10] and to define a version for centrality in graphs [11, 12]. In the context of neuroimaging, graph communicability has been applied to evaluate the contribution of SC topology in generating FC [13]. However, graph theory is often applied in an off-the-shelf manner and this type of approach is limited for explaining the time-series nature of the fMRI measurements.

In this context the present study aims to describe the fMRI-related functional associations between the nodes in the brain network, also known as regions of interest (ROIs), while properly taking time into account. We follow recent works that employed dynamic models of the brain activity to link SC and FC [14]. A great variety of network designs has been explored to combine experimental data in various levels of detail [15, 16, 17, 18, 19, 20]. These dynamic models typically involve a connectivity matrix that describes how the activity propagates in the network. This should be contrasted with another active line of research focusing on the ‘dynamic FC’ that relies on sliding time windows to capture the statistical (functional) dependences between ROIs on the timescale of a minute [21]. The statistical analysis of these successive connectivity measures can lead to the definition of “states” for the whole network or ROIs [22, 23, 24, 25]. Recently, the transitions between the obtained states defined as correlated patterns have been studied using hidden Markov models (HMMs) to generate BOLD activity [26]. A common aspect for the second type of studies is the representation of BOLD signals as independent “static” snapshots corrupted by noise, without considering the transition between successive BOLD activities (akin to time-lagged correlations). Instead, we aim to address this limitation by providing a network-oriented analysis that takes into account the propagating nature of BOLD signals.

Definition of integration measures have been proposed to quantify how nodes in the network exchange information at the scale of the whole network, thus building a global workspace [27, 28]. Previous definitions of integration have focused on the similarity as measured using mutual information or the cross-correlation between the observed activity of subgroups of nodes in the network [29, 30, 31]. The resulting ‘network complexity’ of the network activity reflects the superposed contribution of hubs and network motifs such as modules [32]. Moreover, when network analysis is used with generative models to interpret the collective pattern of interactions between ROIs or quantify integration, it is most often applied on the model activity, such as the model dynamic FC to take time into account [33, 34, 35]. Therefore, it can be argued that these previously-proposed measures for integration focus on the observed or generated activity rather than their causes. To explore this point we will compare network analyses based on the ROI correlation pattern and based on measures of the causal interactions between ROIs in a model.

We present a model-based approach that focuses on the effective connectivity (EC) instead of the FC or SC as a basis for network analysis. The rationale is the following: The dynamic model is an assumption (a prior) about the spatio-temporal structure of the empirical time series and the estimation is a “projection” of the BOLD signals on the space of model parameters. The tuned model can then be used to examine the interactions between the brain regions, via its Green function that quantifies the network response to impulse in given ROIs. Doing so, we incorporate the temporal dimension in the network analysis as EC captures the BOLD dynamics. To illustrate our framework, we use the multivariate Ornstein-Uhlenbeck (MOU) process. Its connectivity, which we term MOU-EC, measures the directional interactions between ROIs in generating the model BOLD activity, while assuming stationarity [18, 36]. The mathematical tractability of the Green function for the MOU process allows for an intuitive interpretation of the MOU-EC in terms of interactions across time between the brain regions [37]. This provides a consistent analysis based on the same network dynamics from the estimation to the interpretation, in contrast to previous formalisms that applied “artificial” dynamics on static networks obtained from neuroimaging data [38, 39, 40, 8, 10]. Although the present study uses MOU-EC, we stress that our framework can be used with other models provided their Green function can be evaluated (e.g. linearized dynamic causal model [41]) or any directed connectivity matrix that could be calculated for brain networks as with Granger causality analysis [42, 43, 44]. The relationship between MOU-EC and other approaches are presented in Methods and further discussed at the end of this paper.

After recapitulating important points about the MOU-EC estimation from BOLD signals [18], we present in Methods the formalism to analyze complex network dynamics with linear coupling [37]. It is based on a core graph-like measure for the interactions between brain regions across time, *“dynamic” communicability*, which serve as the basis for multivariate descriptors. We illustrate the framework using resting-state fMRI and diffusion MRI of the ARCHI database acquired by the European project Connect [45, 46] and available from the Human Brain Project Neuroinformatics platform. In particular, we show how communicability provides a finer and more general description of the network dynamics —by capturing the propagation of the BOLD signals throughout the whole network via EC— than classical analysis based on FC or SC. At the end, we perform community analysis and show how the network displays integration of information, first locally and then globally.

## 2. Methods

### 2.1. Acquisition of resting-state fMRI time series and structural connectomes

The analyses were applied to the ARCHI database [45, 46] composed of 77 subjects (32 females, mean age 23.65 years, with standard deviation 5.16) with high quality T1-weighted images and diffusion data acquired on a Magnetom TimTrio 3T MRI System (Siemens Healthcare, Erlangen, Germany). The MRI protocol included:

- a T1-weighted MRI scan using a MPRAGE pulse sequence with the following settings: field of view FOV = 256 mm, flip angle FA = 9°, inversion time TI = 900 ms, echo time TE = 2.98 ms, repetition time TR = 2300 ms, 160 slices, slice thickness TH = 1.1 mm, in-plane matrix 256 × 256, read bandwidth RBW = 240 Hz/pixel;
- a calibration fieldmap scan using a double gradient echo 2D pulse sequence with the following parameters: FOV = 220 mm, FA = 60°, echo times TE1/TE2 = 4.92/7.38 ms, TR = 500 ms, 35 slices, TH = 3.5 mm, in-plane matrix 64 × 64, RBW = 200 Hz/pixel;
- a resting-state fMRI scan using a 2D gradient-echo echoplanar pulse (GRE EPI) sequence with the following parameters: FOV = 192 mm, FA = 81°, TE = 30 ms, TR = 2400 ms, 40 slices, TH = 3.0 mm, in-plane matrix 64 × 64, RBW = 2442 Hz/pixel, no parallel imaging option, partial Fourier factor = 1.0; each session has 244 frames for a total scan time of 9 min 46 s.

The T1-weighted structural MRI data was processed using Freesurfer 5.3 software [47], which performed cortical tissue (gray and white matters) segmentation, and built surface meshes for both pial interface and grey/white interface of each hemisphere. Surface alignment was performed to an inter-subject template, allowing the projection of the Desikan atlas on these anatomical meshes [48]. The fMRI final activity was obtained by averaging the BOLD signals projected on each mesh vertex over each ROI of the Desikan atlas, see Table 1 below.

**Table 1:** List of the 34 ROI labels for the Desikan parcellation [48]. The order of Fig. 5a, from bottom to top. The parcellation has 1 such ROI for each hemisphere.

| Label | Description |
| --- | --- |
| CUN | cuneus |
| PCAL | pericalcarine gyrus |
| LING | lingual gyrus |
| LOCC | lateral occipital cortex |
| FUS | fusiform gyrus |
| BSTS | bank of superior temporal sulcus |
| IT | inferior temporal cortex |
| MT | middle temporal cortex |
| ST | superior temporal cortex |
| TT | transverse temporal cortex |
| TP | temporal pole |
| PCUN | precuneus |
| SMAR | supramarginal cortex |
| IP | inferior parietal cortex |
| SP | superior parietal gyrus |
| PARC | paracentral gyrus |
| PREC | precentral gyrus |
| PSTC | postcentral gyrus |
| ENT | entorhinal gyrus |
| PARH | parahippocampal cortex |
| INS | insula |
| CAC | caudal anterior cingulate cortex |
| RAC | rostral anterior cingulate cortex |
| PC | posterior cingulate cortex |
| ISTC | isthmus of of cingulate cortex |
| SF | superior frontal cortex |
| CMF | caudal middle frontal cortex |
| RMF | rostral middle frontal cortex |
| POPE | pars opercularis |
| PORB | pars orbitalis |
| PTRI | pars triangularis |
| LOF | lateral orbitrofrontal medial cortex |
| MOF | medial orbitrofrontal medial cortex |
| FP | frontal pole |

Individual SC matrices were built using Diffusion weight imaging (DWI) following the processing steps of the Connectomist-2.0 software [49]: artifact correction, geometrical distortion correction, analytical Q-ball model [50], streamline probabilistic fiber tracking inside an improved T1-based brain mask [51] using 27 seeds per voxel at the T1 image resolution and propagation step size of 0.4 mm. Unreliable short fibers under 30 mm were removed, then region-to-region SC matrices could be built by counting fibers connecting regions of a given atlas [52]. From the individual SC matrices, we calculated a generic SC matrix by averaging over all subjects.

The Constellation parcellation is a refinement from the Freesurfer parcellation based on the DTI data. It subdivides each Freesurfer ROI according to the structural connectivity profile of each ROI point such as to obtain 3 to 5 Constellation ROIs with increased SC homogeneity [52].

Pre-processing of the fMRI dataset included a first step to verify the absence of any outlier (i.e. corrupted volume by spikes, etc.). Standard preprocessing was performed using Nipype [53], including slice timing and motion correction, as in previous studies using the ARCHI dataset [54]. The T1-weighted scan was used as a reference and both fMRI and fieldmap datasets were matched to this reference using a mutual-information based registration algorithm providing the corresponding rigid transformations T(fMRI→T1) and T(B0→T1). Unwrapping of the phase difference map included in the fieldmap scan was performed using a weighted full multigrid phase unwrapper algorithm that did not require the preliminary definition of a mask of the brain, and the unwrapped phase map was then matched to the fMRI dataset by composition of the former transformations in order to correct for susceptibility artifacts. Finally, bandpass filtering with [0.01 – 0.1] Hz was applied to the signal resulting from the averaging over each ROI.

### 2.2. MOU process fitted on whole-brain fMRI data

A motivation for using the MOU process is because its Green function has an analytical formulation, related to the matrix exponential [37]. The MOU process is determined by (i) a local leakage related to the decay time constant *τ*, (ii) a directed weighted matrix A associated with linear coupling that is the MOU-EC and (iii) fluctuating inputs with zero mean and a given covariance matrix Σ. Assuming stationarity over each resting-state session, we estimate *τ, A* and Σ using a form of gradient descent performed without simulating the network activity, but using analytical consistency equations. This optimization procedure has a single solution in the absence of observation noise [18].

The following mathematical details will be useful for the calculation of our graph-like measures. Formally, the MOU process in matrix form is defined as

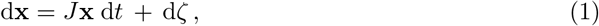

where the Jacobian matrix *J* is determined by the decay time constant *τ* (identical for all ROIs) and *A_ij_* is the MOU-EC weight from ROI *j* to ROI *i* (different from the usual convention in graph theory):

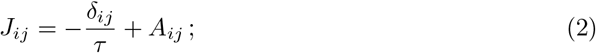

here *δ_ij_* is the Kronecker delta (equal to 1 when *i* = *j* and 0 otherwise). Similar to a transition matrix for a Markov chain, the matrix *A* determines the Jacobian of the dynamic system. The time constant *τ* is an abstraction of the hemodynamic response decay. The choice of a single time constant *τ* for all ROIs, but individual for each subject, comes from the data, which exhibit a smaller variability across ROIs than across subjects (not shown here). Last, the fluctuating input to ROI *i* is denoted by *ζ_i_* and represents spontaneous activity. It is modeled as a Wiener process (temporally white noise) with covariance matrix Σ. Here Σ is kept diagonal, meaning that *ζ_i_* are independent white noise, but in general it may also involve cross-covariances [36]. To ensure stable dynamics, the Jacobian *J* must have eigenvalues with strictly negative real part. In particular, the dominating eigenvalue (or spectral diameter) of the weight matrix A needs to satisfy λ_max_ < −1/*τ*, meaning that the local leakage must be sufficiently quick (or strong) compared to the global network feedback.

The MOU-EC weights *A_ij_* come from the tuning of the model to reproduce the spatio-temporal structure of the BOLD signals, which is simply the pair of covariance matrices FC0 and FC1 in Fig. 3a, without and with a time lag equal to 1 TR (temporal resolution of the BOLD measurements). The Lyapunov optimization (akin to a gradient descent) iteratively adjusts the weights *A_ij_* for each existing connection, the variance Σ_*ii*_ for each ROI and a common *τ* for all ROIs to reproduce the covariances of fMRI signals involving time shifts (empirical FC matrices). The optimization steps are repeated until reaching a minimum in the model error, which is the matrix distance between corresponding model and empirical FC0/FC1. Details can be found in previous papers [18, 36]. Note that we use simplified notation for compactness here, as compared to those previous publications.

The estimated MOU-EC is in general a directed and weighted matrix [18]. EC is also “sparse” with density 34%, corresponding to 1562 non-zero elements among the *N*(*N* − 1) = 4556 off-diagonal matrix elements. Its topology —which interregional connections exist in the model— is determined by thresholding the average SC matrix for all subjects described in the previous section. This reduces the number of parameters (including 68 for Σ) and improves the robustness of the estimation as each session has 244 × 68 = 16592 time points over all ROIs. Importantly, the individual values of the estimated MOU-EC weights do not depend on SC values, but are determined by the optimization procedure for the model FC that best reproduces the empirical FC.

The model optimization approach contrasts to previous studies that used SC values for the connectivity weights in their model, while tuning local dynamics such as Kuramoto oscillators or mean-field approximation of spiking neurons [55, 56, 17]. Our estimation procedure can be seen as a “projection” of the empirical spatio-temporal BOLD structure on the space of model parameters, where MOU-EC captures the propagating nature of the BOLD signals.

We use the EC terminology that originates from electrophysiology and was developed by the literature of dynamic causal model. In our case MOU-EC is the underlying connectivity in a generative dynamic model of BOLD activity, which corresponds to a historical key aspect of the EC concept. However, our model does not involve the hemodynamic response and differs from state-space models such as the dynamic causal model that separates the neural network and the generation of BOLD activity [15, 57]. The limitations of MOU-EC estimation compared to other methods will be reviewed in the end of the Discussion.

### 2.3. Dynamic communicability to measure interactions between ROIs across time

Following our previous paper [37], we define *dynamic communicability* to characterize the network interactions induced by its weighted and directed connectivity *A*, ignoring the input properties Σ. Our definition is adapted to study complex networks associated with realistic (stable) dynamics where time has a natural and concrete meaning, with the Jacobian *J* combining *A* and the leakage time constant *τ*. In comparison, a previous version of communicability for graphs [8] relied on abstract dynamics. From the graph communicability *e^A^* based on the matrix exponential, the strengths Σ_*i*_(*e^A^*)_*ij*_ and Σ_*j*_(*e^A^*)_*ij*_ were used to evaluate the roles for nodes as broadcasters or listeners in the network [10]. The diagonal elements (*e_A_*)_*ii*_ quantify the feedback received by a node from its own abstract “activity” and was used to define centrality [11]. Getting inspiration from those studies, we base our framework of dynamic communicability on the Green function (or network impulse response) to perturbation for the MOU process, focusing on the temporal evolution of such interactions (see dashed arrow in Fig. 2a). The present study uses the MOU process because of its analytical tractability, but our framework can be applied to any local dynamics for which the Green function is known.

Dynamic communicability measures the network response when applying a perturbation to a given node (node 1 in Fig. 2a). Importantly, communicability excludes the “intrinsic” relaxing response due to the local leakage to focus on the contribution of the connectivity *A* to the network response, see the zero communicability for node 1. Note that the strong weight from node *A*_23_ from node 2 to node 3 gives to a strong communicability between 1 and 3, even though *A*_21_ is small. For recurrent networks, the existence of feedback loops gives a positive communicability for a node onto itself; see the green and blue curves in the top and middle configurations in Fig. 2b. For the MOU process, communicability is the “deformation” of its Green function *e^Jt^* due to the presence of the matrix A (dashed curves), as compared to the Green function *e*^*J*^0^ *t*^ corresponding to the Jacobian with leakage only and no connectivity, 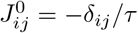 (dotted curves). It corresponds to the family of time-dependent matrices (see also Fig. 3a in Results):

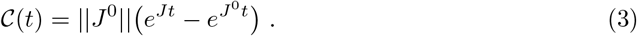

Recall that *t* ≥ 0 here is the time for the propagation of activity in the network, referred to as ‘integration time’, which is different from the time in Eq. (1). The scaling factor 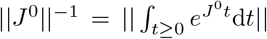 where || · || is the L1-norm for matrices (i.e. sum of elements in absolute value) is used for normalization purpose, e.g. when comparing networks of distinct sizes [37].

From each matrix 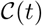, we define the total communicability that sums all interactions

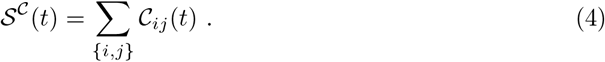

Total communicability measures the effect of a common perturbation at all nodes after the integration time *τ*, reflecting the global network feedback determined by the matrix *A*. It relates to eigencentrality as the function 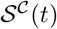 is mostly determined by the largest eigenvalue of *A* [37]. We also define the diversity (or heterogeneity) among the ROI interactions in the time-dependent matrices 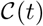, which can be seen as a proxy for their homogenization over time:

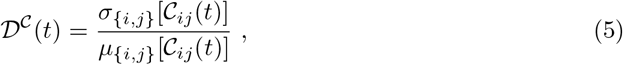

defined as a coefficient of variation where *μ*_{*i,j*}_ and *σ*_{*i,j*}_ are the mean and standard deviation over the matrix elements indexed by (*i,j*). In Fig. 2b the top configuration has the same communicability from node 1 to node 2 (in purple) as from 2 to 1 (in cyan), because the corresponding weights in *A* are identical. For the unbalanced connectivity in the bottom configuration, the communicability from 1 to 2 is larger than the converse. Because both network configurations have the same sums *A*_12_ + *A*_21_, they have very similar total communicability 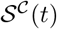 in Fig. 2c, but distinct diversity 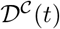.

### 2.4. Community detection

To detect communities based on communicability, we rely on Newman’s greedy algorithm for modularity [58] that was originally designed for weight-based communities in a graph. Adapting it here to the matrix 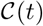 at a given time *t*, we seek for communities where ROIs have strong bidirectional interactions. In the same manner as with weighted modularity, we calculate a null model for EC:

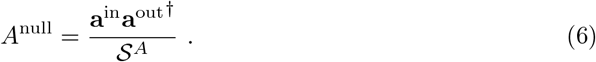

Note that we preserve the empty diagonal. The resulting matrix contains from the expected weight for each connection, given the observed input strengths **a**^in^ and output strengths **a**^out^; 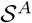 is the total sum of the weights in *A*. Then we caclulate 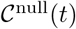 using Eq. (3) with *A*^nu11^ instead of *A*. Starting from a partition where each ROI is a singleton community, the algorithm iteratively aggregates ROIs to form a partition of *K* communities denoted by *S_k_* that maximizes the following quality function:

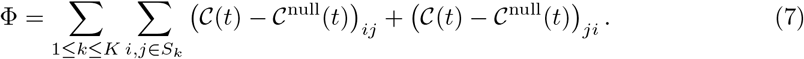

At each step of the greedy algorithm, the merging of two of the current communities that maximizes the increase of Φ is performed.

## 3. Results

This section uses our definitions of dynamic communicability to interpret resting-state fMRI data (see Fig. 3a). The data were recorded from 77 subjects during 244 frames each separated by a TR = 2.4 s with eyes closed in a scanner. Our framework relies on a dynamic system and, for illustration purpose, the present study uses our recent optimization to fit the MOU model to BOLD signals [18, 36]; the python code is available on github.com/MatthieuGilson/WBLEC_toolbox). This model optimization combines several key aspects that are developed in Methods:

- A whole-brain approach [16] is necessary to properly take into account the distributed nature of information conveyed by BOLD signals, as was experimentally observed for high-level cognition [59] or neuropathologies [60].
- The MOU process is optimized to reproduce the spatio-temporal structure of the BOLD signals, which was shown to convey information about the behavioral condition of subjects [61, 62, 63]. This contrasts to traditional analyses of the “spatial” FC, which does not involve time lags.
- The optimization combines information from both FC and SC data in estimating the MOU-EC [18], which is then subject/condition-dependent with a “sparse” topology corresponding to the anatomy.
- In the model the directional connections between brain regions describe causal interactions between ROIs. However, the hemodynamic response is not explicitly modeled in our MOU process, which will be discussed later.

The use of whole-brain directional connectivity is the feature that makes the application of dynamic communicability interesting. Another motivation fo use of MOU-EC is that the leakage time constant *τ* that is part of the Jacobian *J* in Eq. (2) is jointly estimated during the model optimization. In contrast, taking SC or another directed connectivity matrix (e.g. obtained using Granger causality) as matrix *A* in *J* to use Eq. (3) would require an arbitrary choice for *τ*, because they are not estimated with the dynamic model. Note also that the heterogeneity of the Σ estimates (see purple arrows with various thicknesses in Fig. 3a) is ignored in the present analysis and left for future work.

We firstly show how communicability provides information about the topology of the estimated MOU-EC (leaving out the estimated input variances Σ), from the global level of the whole network to the local level of individual connections or ROIs, including the detection of communities of ROIs.

### 3.1. Communicability provides both global and local information about the brain network

In the present formalism, communicability is the family of matrices 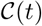 in Fig. 3a that describe the interactions between pairs of ROIs across time. In each matrix 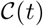, a column measures how a unit perturbation applied at *t* = 0 to the corresponding source ROI propagates throughout the network via the recurrent MOU-EC and impacts the other target ROIs after a delay *t* (termed ‘integration time’) during which the network effect builds up. The theory is based on the Green function or impulse response of the network, see Eq. (3) in Methods. This directed measure thus integrates all possible pathways between the source and target ROIs, while taking the nodal dynamics of the model into account (here a exponential decay related to a time constant *τ*, see Fig. 1c). For the resting-state, perturbations are the fluctuations at each ROI, whose effects sums over time.

**Figure 1:**
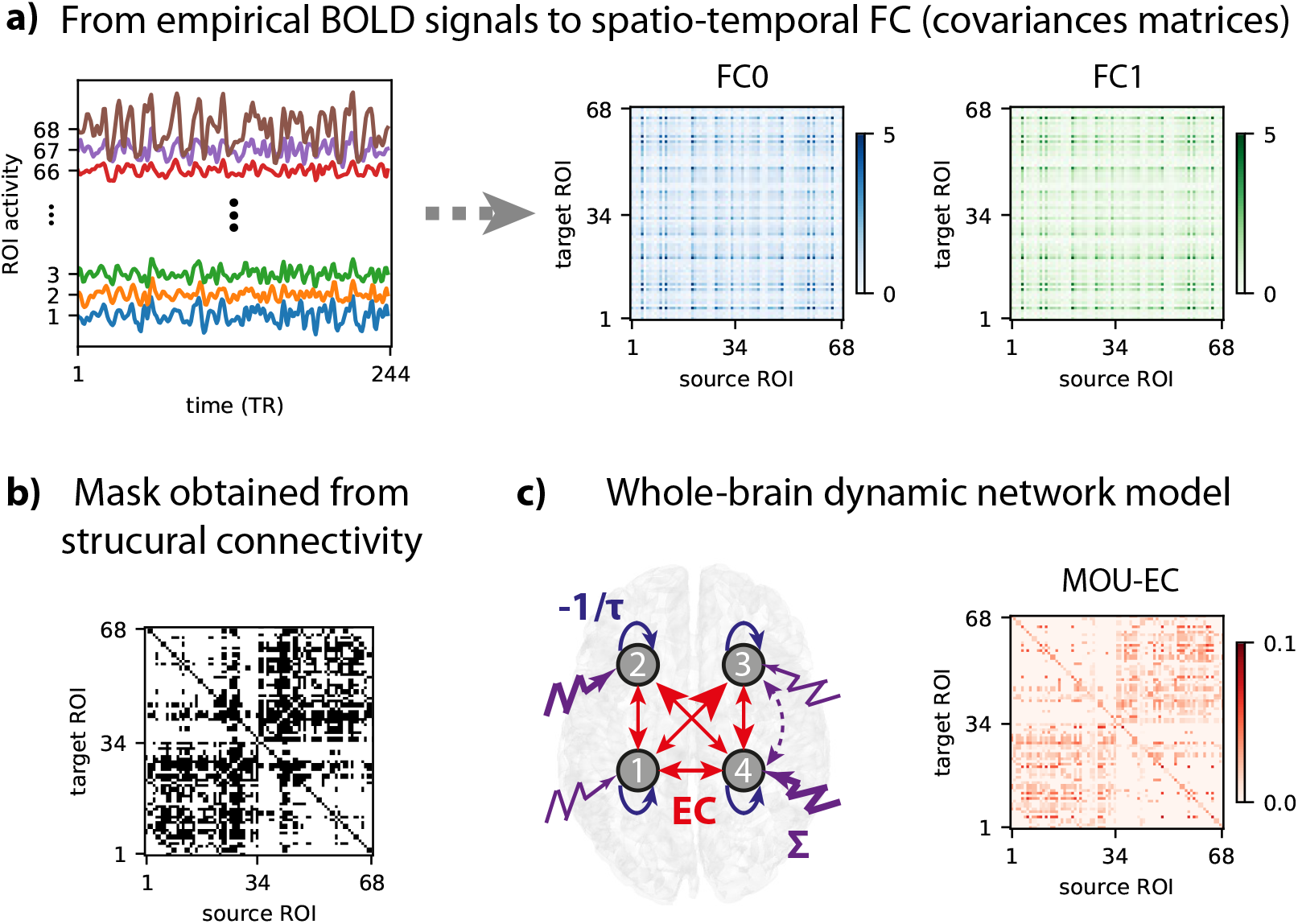
MOU-EC estimation. **a)** From the fMRI signals (left plot), the covariance matrices FC0 and FC1 are calculated, represented in blue and green respectively. FC0 is the zero-lag covariance and FC1 corresponds to a time shift of 1 TR. **b)** Topological mask for existing connections in the model obtained from thresholding the average SC over subjects. **c)** The network parameters comprise of the MOU-EC matrix A (red arrows in the left diagram) and self-inhibition corresponding to the time constant *τ* (blue arrows), see Eq. (2). Note that *A* is not a full matrix, because its topology is determined by the mask obtained from SC in panel b. In addition, the local excitabilities or inputs (purple arrows) are determined by their covariance matrix Σ. Here only 4 ROIs are represented for readability. In this paper the focus in on the estimated MOU-EC matrix, corresponding to matrix *A* in Methods, as well as the leakage time constant *τ* involved in Eq.(2). For each session, *A* is estimated from the covariance matrices FC0 and FC1, capturing the average BOLD transition statistics over the session (assuming stationarity). Further details about the dynamic model and estimation procedure can be found in previous studies [18, 36].

**Figure 2:**
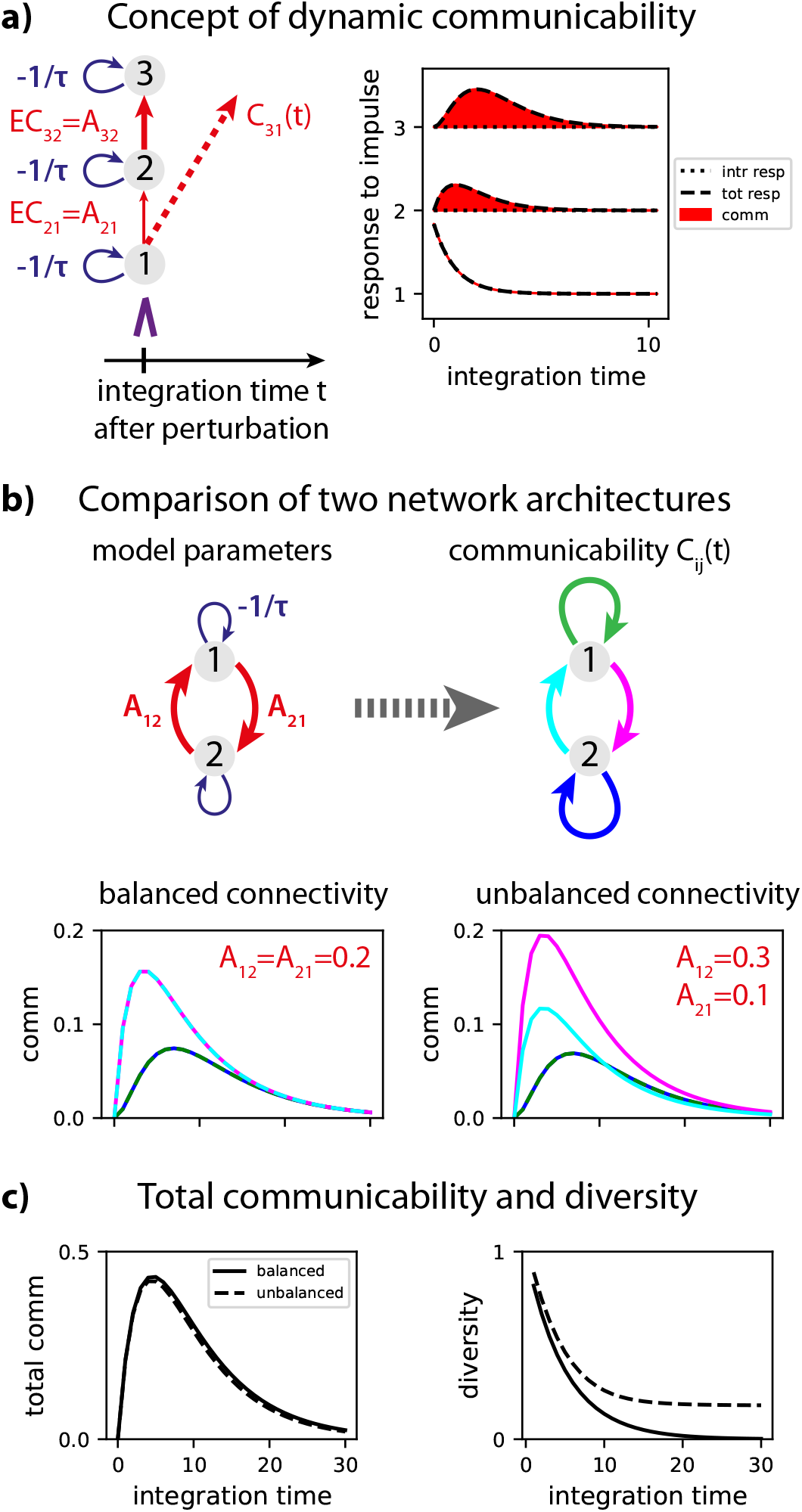
Concept of dynamic communicability. **a)** Communicability for three nodes relies on the response of the network to a unit perturbation at a given node (here node 1). The dynamics are determined by local leakage related to the time constant *τ* and the connectivity weights *A* (left diagram). In the right plot the total response of the is represented in dashed line and the “intrinsic” response due to leakage only in dotted line, corresponding to *J*^0^ in Eq. (3). The communicability 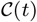 is the difference between these two curves. **b)** The left diagram represents a network of two nodes with recurrent connections. The communicability 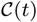 is a 2 × 2 matrix with 4 elements (see green, blue, purple and cyan arrows in the right diagram). The bottom plots comparison the communicability for two network configurations. Note that the blue and green curves are superposed. **c)** Total communicability 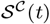 in Eq. (4) and diversity 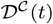 in Eq. (5) for both networks. The solid curves correspond to the balance network and the dashed curves to the unbalanced one.

Initially aligned with MOU-EC (i.e. interactions through the strong direct connections dominate), the pattern of communicability progressively reshapes and the superiority of connections with strong weights dilutes, as illustrated in Fig. 3b. In particular, unconnected regions may have strong communicability due to network effects (see crosses for *A_ij_* = 0 on the left of the plots). This homogenization results from the superposed loops in the MOU-EC matrix that generate a strong overall feedback that distributes the effect of the fluctuations.

**Figure 3:**
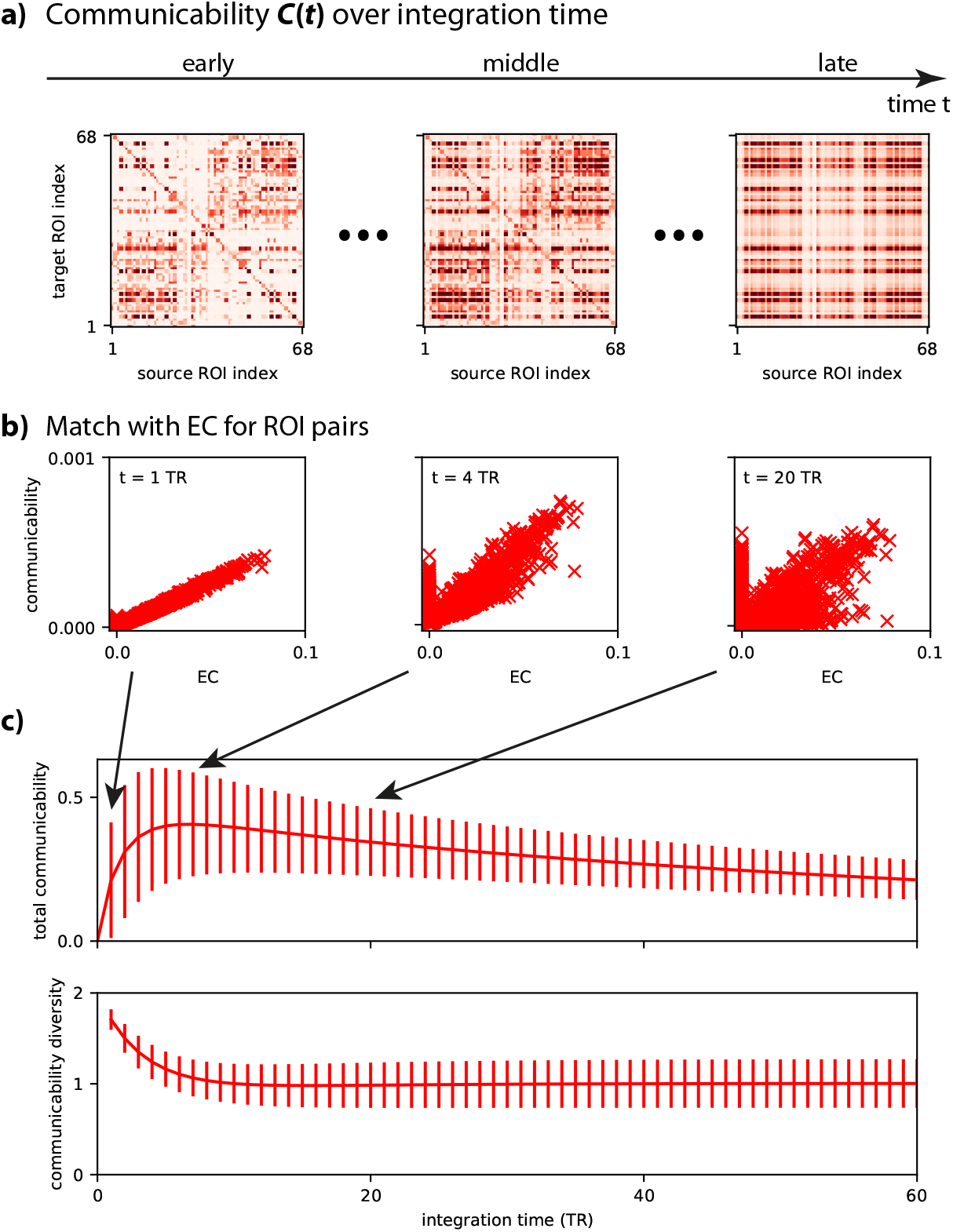
Whole-brain communicability. **a)** Communicability for a single subject, corresponding to the family of matrices 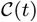 in Eq. (3) that involves the matrix exponential of the Jacobian multiplied by time *t* ≥ 0. Recall that t corresponds to the integration time for fluctuating activity in the network. The range is the same as the MOU-EC matrix in panel a. **b)** Match between MOU-EC and communicability for each pair of ROIs (red crosses) at three times for the same subject as in panel b. **c)** Total communicability 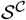 defined as in Eq. (4) reflects the global network feedback. The x-axis represents the integration time in units of the scanner measurements (time resolution, or TR, equal to 2 s). The curves correspond to the average over the 77 subjects and the error bars the standard deviation. Diversity of communicability 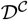 defined as in Eq. (5) that quantifies the homogenization of interactions across time.

The strong network feedback is reflected in the total dynamic communicability 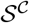 in the top panel of Fig. 3c, which rises to reach a maximum around 7 TR, before decaying very slowly. The integration time of the peak corresponds to the lag after which a perturbation in the network (here affecting all nodes) has the maximum influence (given by the size of the peak), due to the building up of the network effect. The slow decay afterwards indicates that the effect persists over time. The homogenization is reflected in the communicability diversity 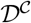 (bottom panel) that quickly decays to stabilize after 15 TR. This defines the temporal horizon after which the effect of perturbations are most broadly distributed across the network. Note that, meanwhile, 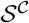 is still high. These curves provide a signature for the estimated MOU-EC, which can be analyzed to uncover its topological properties.

### 3.2. Comparison with reference networks

Communicability enables the quantitative comparison of the estimate for brain dynamics with other reference network models. Here we compare the global measures of total communicability 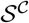 and its diversity 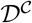 in Fig. 3d to their counterparts for surrogate networks obtained by manipulating the MOU-EC estimates. The comparison between the timescales associated with the corresponding curves provides insight about equivalent topologies to generate dynamics similar to the model estimates. Topological properties such as strong feedback and short path length are reflected in 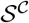 and 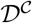 [37].

In Fig. 4d, the evolution of 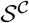 and 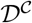 for the original data (red curves) is compared with four surrogate models:

- The random surrogate in Fig. 4a reallocates the weights to a pair of ROIs. The exact weight distribution is conserved in the network, but not for each ROIs.
- The null model in Fig. 4b was proposed for Newman modularity [58] to detect communities and corresponds to resulting a full matrix with for each pair of ROIs the expected weight that preserves the distribution of input and output strengths over each ROI, see Eq. (6) for details.
- The ring surrogate in Fig. 4c (left matrix) reorganizes the input connections for each ROI to promote local connectivity with a ring topology determined by the index of the ROIs. In this arbitrary topology, distant ROIs are not directly connected.
- The shortened ring in Fig. 4c (right matrix) is similar to the previous ring surrogate, but with pooling every 3 input connections, resulting in lower density (by a factor 3).

**Figure 4:**
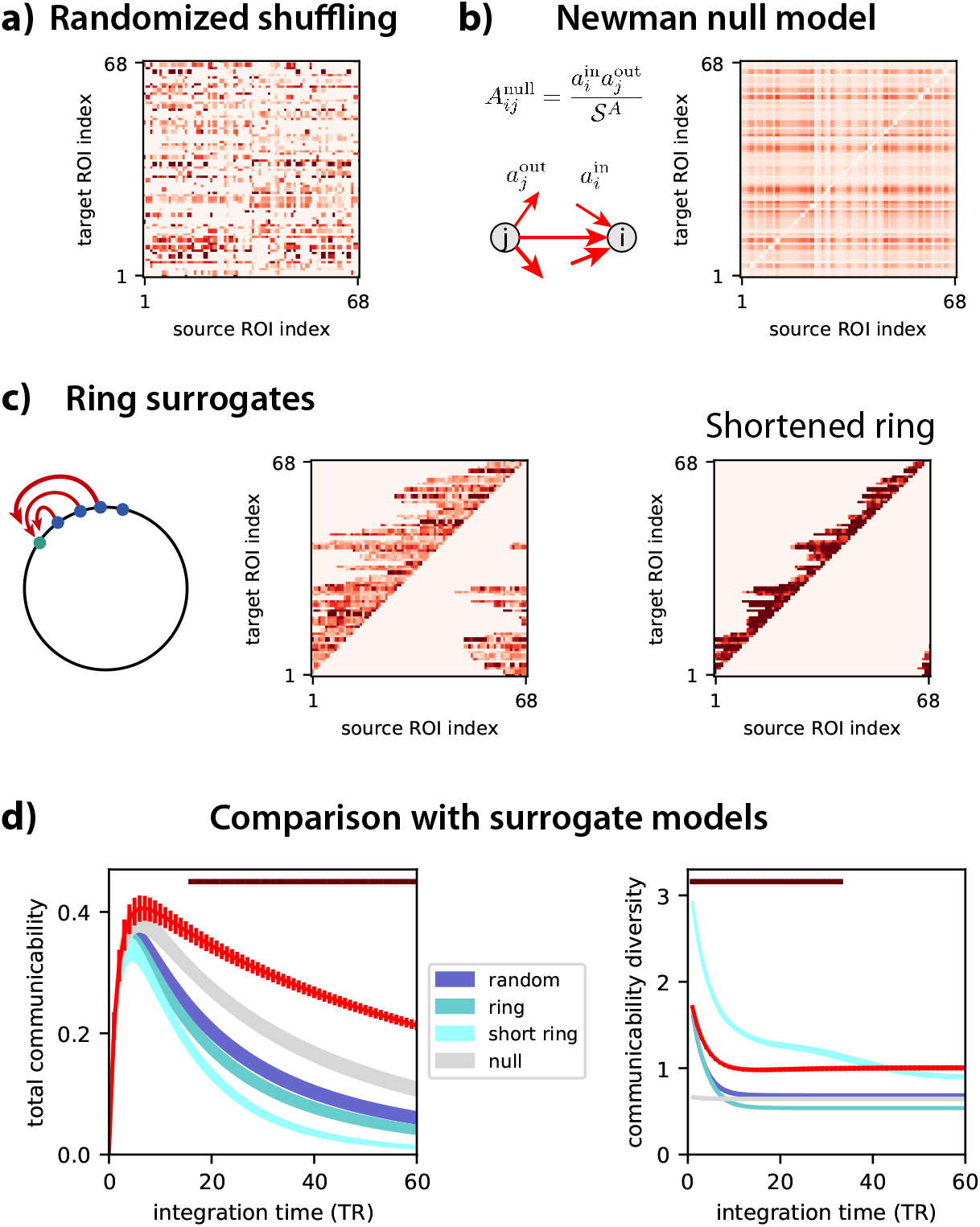
Comparison of the estimated model with surrogates. **a)** Randomized version of the estimated matrix *A* by globally shuffling the target ROIs. The range of the matrix (as well as others in this figure) is the same as in Fig. 3b. **b)** The null-model connectivity is calculated using Eq. (6) redistributes the weights while preserving the input and output strengths for each ROI. This results in a full matrix, apart from the empty diagonal. **c)** The ring surrogates consider the initial ROI order and reallocate the non-zero weights in the original *A* to the preceding ROIs. The ring preserves the number of connections (hence the density), whereas the shortened ring corresponds to grouping connections three by three, resulting in fewer connections. **d)** The total communicability 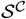 and the corresponding diversity 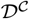 in red are the same as in Fig. 3d, but the error bars indicate the standard error of the mean here (much smaller than the standard deviation). The colored areas correspond to the surrogate networks in panels a to c, which preserve part of the connectivity statistics of the original *A* (see Table 2). The transformations being applied individually for each subject. The variability of the curves (represented by the thickness of the plotted area) corresponds to the standard error of the means over subjects, as error bars for the red curve. The dark red bars over the curves indicate the integration times where the model is significantly different from all surrogates, as measured by the Mann-Whitney test for which the p-value is smaller than 0.001 for all 4 comparisons (uncorrected).

In each case, the shuffling of the original estimate is performed for each subject and the average over subjects is then calculated. Importantly, these surrogates all destroy parts of the statistics of the original weight distribution in the estimated *A* in a specific manner while preserving others. They all preserve the total weight in *A*, which is the 1st-order moment of the weight distribution, and leave the diagonal empty. For example, the random and ring surrogates both preserve the connectivity density (1st-order of the overall distribution for the binarized weight matrix) as they topologically reallocate weights from connection to connection in a one-by-one mapping, but only the ring surrogate preserves the original sum of input weights for each ROI. This is summarized in Table 2, where the mean input and output weight per ROI, which is the first order of the weight distribution “projected” in one dimension of the weight matrix. Note that each surrogate also destroys the second-order statistics, corresponding to the joint distribution of weights for pairs of ROIs.

**Table 2:** This table shows which properties of the original network are preserved (✓) or randomized (✔) by the surrogates.

| Property | random | null | ring | short ring |
| --- | --- | --- | --- | --- |
| Mean weight | ✓ | ✓ | ✓ | ✓ |
| In weight for each ROI | ✗ | ✓ | ✓ | ✓ |
| Out weight for each ROI | ✗ | ✓ | ✗ | ✗ |
| Density | ✓ | ✗ | ✓ | ✗ |

The random surrogate (dark blue curve) exhibits a smaller values for 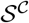 and a homogenization slightly after 20 TR, but at a lower asymptotic value for 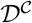. The ring surrogate (dark cyan curve) destroys the local loops and clusters, which decreases 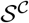 even more strongly than for the random surrogate. In our previous theoretical study, we used ring lattice to study how communicability captures the small-worldness in networks, which corresponds to a quick stabilization of 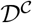 compared to the timescale of 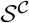 [37, Fig. 6]. Here the estimated MOU-EC exhibits a profile for 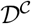 that is much closer to the random surrogate than the ring lattice, indicating that the fluctuating activity quickly propagates throughout the whole network.

Because both surrogates have the same density and individual weights, this suggests a specific internal organization for the estimated EC, related to the multiple loops and resulting in a strong network effect.

For the short ring surrogate (light cyan curve), the main difference concerns the much slower homogenization for 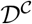. This arises because shortest paths between ROIs are much longer for the short ring (light cyan curve) than the ring surrogate (dark cyan curve). This shows that the mean weight per ROI is not the relevant feature to understand the propagation of activity in the network.

Last, the null model (gray curve) also has a weaker 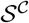, but more interesting is the flat curve for 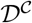. This illustrates the already-homogenized (fully-connected) network for the surrogate obtained by randomizing the second order statistics of the connectivity. This property is the reason for its use as a baseline when performing community detection based on communicability.

To recapitulate, the comparison of those descriptors for the estimated MOU-EC with the surrogates —each specifically destroying part of the weight structure— gives insight about how close or distant the brain network dynamics are with respect to the corresponding stereotypical models. The dark red bars in Fig. 4d indicate statistical difference between the model (in red) and the 4 surrogates with p-value < 0.001 (uncorrected). For 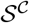, it concerns late integration times when the global network feedback builds up. For 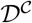, the homogenization saturates at a high value during early integration times (and faster than the short ring surrogate), suggesting a distinct detailed structure of the estimated MOU-EC as compared to the surrogates. A possible explanation is the existence of modules [37, Fig. 7], which will be further studied later using community detection.

### 3.3. ROI-specific analysis and comparison with SC and FC

The brain network structure determines a hierarchy among ROIs. For binary graphs, the notions of degree and centrality have been used to detect highly connected ROIs, or hubs. This type of approach has been used to explore the importance of ROIs in the brain based on SC or FC data [6, 7, 1]. A limitation is that SC and FC do not have directional information (i.e. they are symmetric matrices). Graph communicability [10] can be used to describe differentiated input and output properties for the ROIs. We now show how our dynamic communicability can be used to characterize ROIs in the network and incorporating the temporal dimension.

The overall picture obtained from MOU-EC is differentiated roles across ROIs, each with its own amplitude (in red in Fig. 5a) and timescale (Fig. 5b). In general, ROIs have distinct input and output communicability (Fig. 5c), thus defining listeners and broadcasters. This relates to the propagating nature of fMRI signals, which is related to the asymmetry of estimated MOU-EC matrices [18] and is in line with previous results of the fMRI lag structure [63]. Compared to our previous analyses that focused on individual MOU-EC connections [36, 64], communicability provides a description of the ROI role over time. ROIs usually classified as hubs according to their high degree in SC (in black in Fig. 5a) are indicated by the black crosses: the precuneus (PCUN), superior frontal cortex (SF) and superior parietal cortex (SP). PCUN and SP exhibit strong input communicability, which classifies them as listening ROIs, integrating the activity from the rest of the brain. This is in line with previous results that suggested an global integration role for PCUN as part of the default mode network [65]. In comparison, SF has more balanced (and weaker) input and output communicabilities. The visual ROIs (CUN, PCAL, LING and LOCC) are both listeners and broadcasters. Other noteworthy ROIs with strong output communicability are the pre- and postcentral ROIs (PREC and PSTC), as well as the superior temporal ROIs (ST).

**Figure 5:**
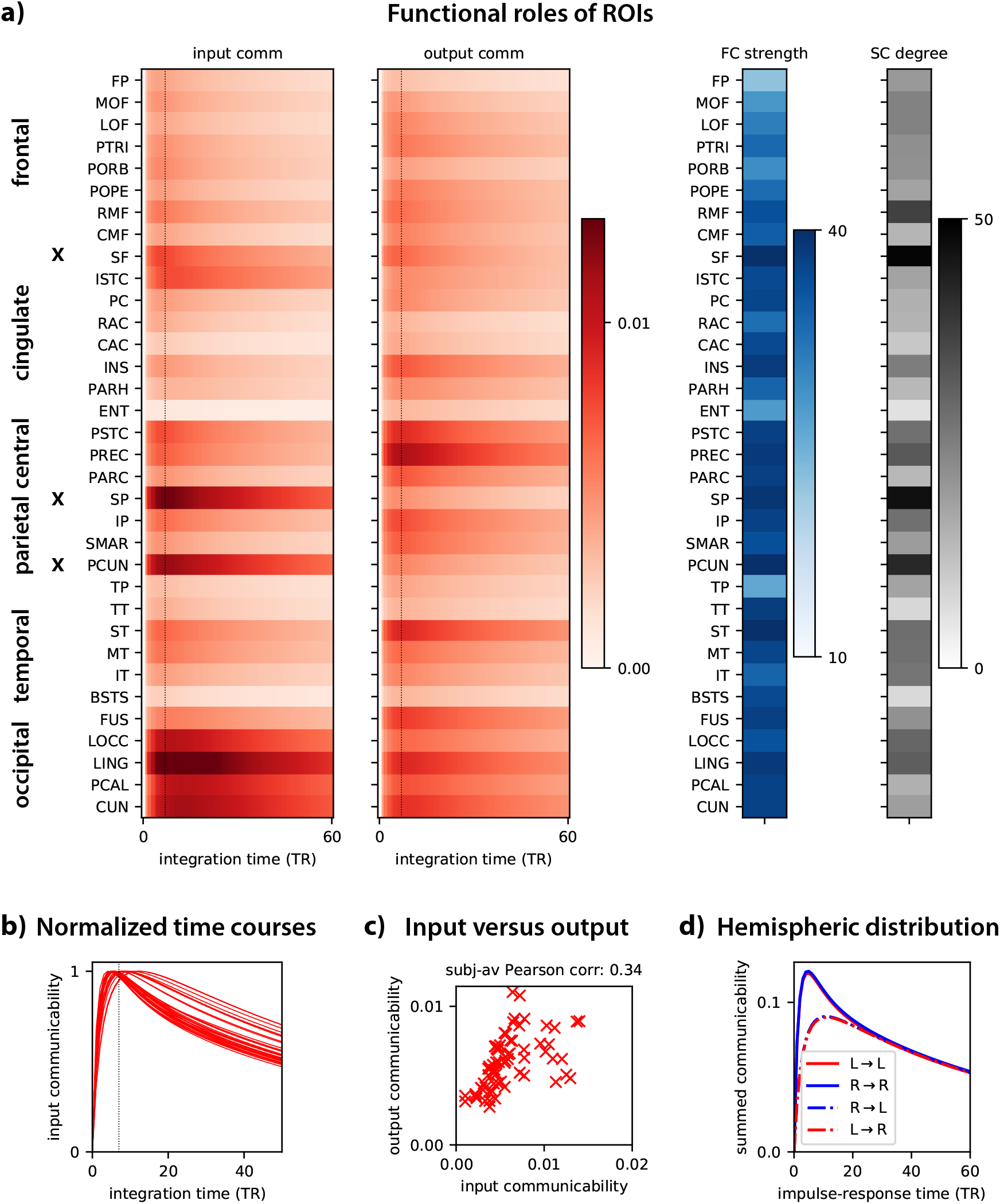
Characterization of ROI roles. **a)** Temporal evolution of input and output communicability for all ROIs (in red). The values are averages over the two hemispheres and all subjects. The ROIs are grouped by anatomical areas and black crosses indicate the three pairs of rich-club ROIs. The dotted vertical lines indicate the maximum for the average 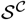 over subjects in Fig. 3d. The right plots indicates the FC strengths (in blue) and SC degrees (in black). Here recall that FC is the classical Pearson correlation between the BOLD signals (not covariances as in Fig. 1). **b)** The normalized time courses for input communicability reveal a variety of timescales across the ROIs. **c)** Comparison of input and output communicability for each ROI (red cross). The plotted value is taken at the maximum *t* = 7 TR, indicated by the dashed vertical lines in panel a. **d)** Evolution of the total communicability within and between the two hemispheres (L for left and R for right).

At the global level, previous work [66] showed asymmetries in the left and right within-hemisphere FC patterns. In contrasts, we observe for our data a symmetric communicability for the two hemispheres in Fig. 5d. Moreover, the communicability is initially higher within each hemisphere (solid curves) than between hemispheres (dashed curves), but rapidly reaches similar values around 20 TR. This points to slightly weaker inter-hemispheric communication compared to within-hemispheric communication at rest. It can be explained by the difference in connectivity degree of the between/within hemispheric MOU-EC weights, as similar curves can be obtained using the binarized MOU-EC estimate with ad-hoc rescaling (giving the matrix in Fig. 1b). Our previous results based on MOU-EC indicated stronger inter-hemispheric communication when analyzing changes of individual MOU-EC weights when engaging a passive-visual task [36], which would be interesting to quantify at the network level using the present formalism.

Comparing FC strengths and SC degrees with communicability in Fig. 5a, we observe that the correspondence between strong values is not trivial. To further quantify this relationship, we display these ROI-specific measures as scatter plots in Fig. 6a-b, where communicability is taken at the peak at *t* = 7 TR (dotted line in Fig. 6a-b). We find that input communicability correlates with the SC degree (dark red dots), as measured by a Pearson correlation of 0.58 (evaluated for each subject, then averaged over them). This suggests that regions with strong input communicability have sufficiently many connections to support it. More moderately, output communicability correlates with FC strengths (purple triangles) with a Pearson correlation of 0.51. It is smaller than 0.35 otherwise. In comparison, the relationship between SC and FC in Fig. 6c shows a stronger correlation (Pearson correlation of 0.66). Even though the general relationship is positive correlations between these descriptors, our model-based approach provides distinct information from the classical analyses based on SC/FC alone. When varying the integration time for input/output communicability, we see that the Pearson correlation is stable in Fig. 6d, except for a transient peak for input communicability and SC degree. This can be explained because of the equal sparsity between these two matrices for early integration times.

**Figure 6:**
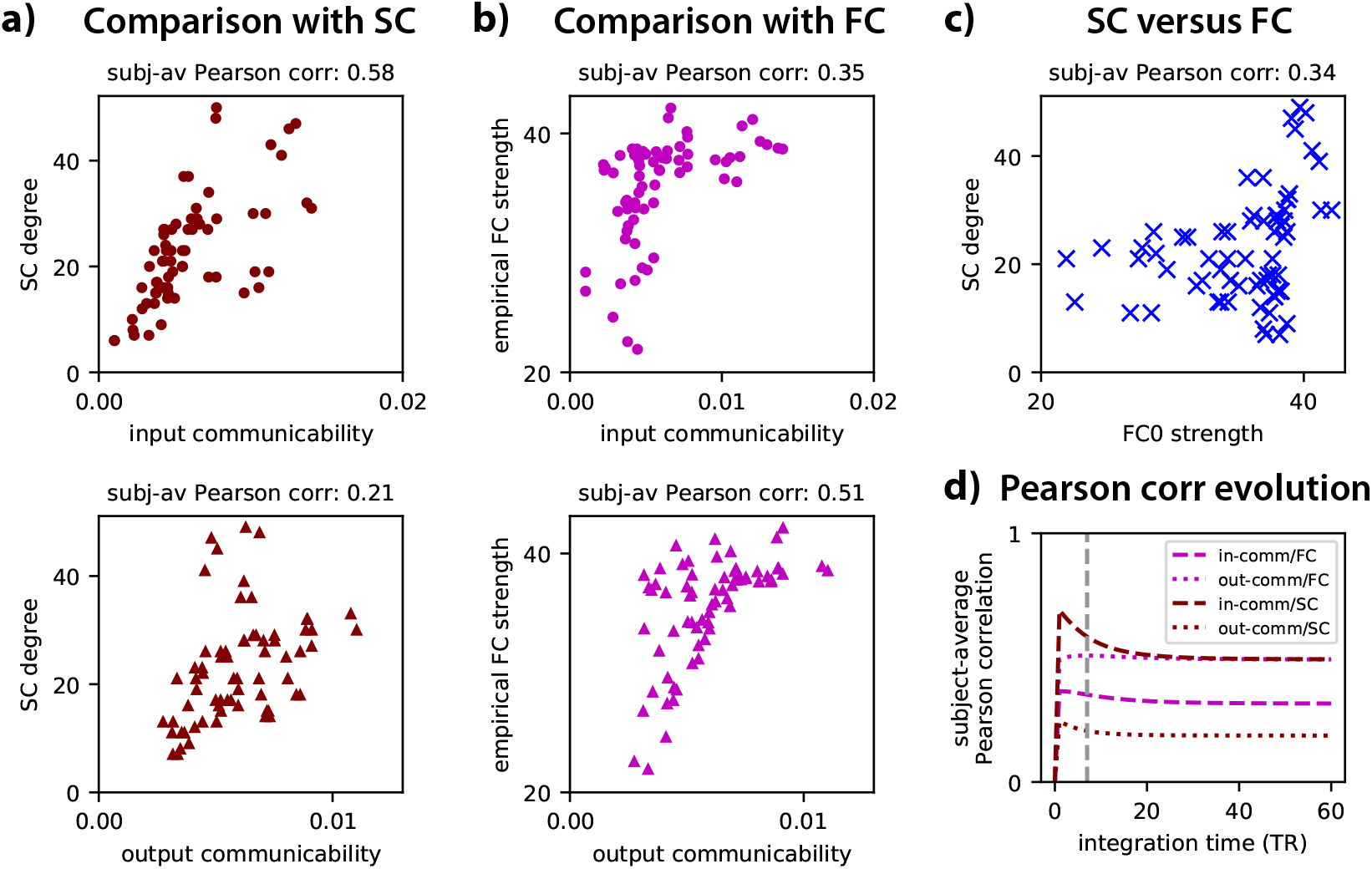
Comparison with information obtained from SC and FC. **a)** Comparison of the input and output communicability (left and right panels in each row with dots and triangles, resp.) with the SC degree. Each symbol represents the value for a ROI, averaged over all subjects; for communicability, it is taken at integration time 7 TR corresponding to the dashed vertical lines in Fig. 5a. The Pearson correlation coefficient above each plot corresponds to the average over all subjects. **b)** Same as panel a for empirical FC strengths. **c)** Comparison between SC degrees and strengths with empirical FC strengths. The value above the plot is the average over subjects of the Pearson correlation between individual input and output communicability. d) Evolution of the Pearson statistics when varying the integration time (x-axis).

Previous studies showed how MOU-EC weights are modulated in a task-dependent manner [36, 64]. In particular, the output MOU-EC weights of hubs are down-regulated at rest as compared to visual tasks [64]. This suggests that not all existing MOU-EC pathways are used at rest, which may explain the very weak correlation of output communicability with SC in Fig. 6.

### 3.4. Functional communities in brain network and information integration

The high saturating value of 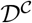 in Fig. 3d is also reminiscent of hierarchical networks consisting of several modules that have rather strong coupling between them, for which the diversity stabilizes early when the total communicability is still large and close to its peak [37, see Fig. 7 there]. To further examine this aspect, we perform a community analysis to reveal the communication between ROI groups over time. In essence, communicability is compared to that obtained with the null connectivity model in gray in Fig. 4. ROIs with greater reciprocal communicability compared to the null model are grouped into communities, yielding the coparticipation matrices in Fig. 7a. These matrices measure the robustness of the communities across subjects: Well-defined correspond to dark squares on the diagonal (after reordering the ROIs), whereas weak associations of ROIs appear in lighter gray. The 4 communities correspond to observed of resting-state networks [67, 68]:

- insularo-temporal and lateral frontal ROIs: CMF, RMF, POPE, PTRI, SMAR, IP, BSTS, ST, MT, TT, INS;
- visual ROIs, precuneus and medial ROIs close to the hippocampus: CUN, PCAL, LING, LOCC, FUS, PARH, ENT, ISTC, PCUN;
- lower frontal and temporal ROIs plus a cingulate ROI: LOF, MOF, PORB, RAC, FP, IT, TP;
- sensorimotor ROIs and ROIs from the default-mode network: PREC, PARC, PSTC, CAC, PC, SF, SP;

in the order of the diagonal blocks from the bottom left to the top right in Fig. 7a with *t* = 1 TR. All communities are symmetric with the two homotopic ROIs in both hemispheres. The full name regions can be found in Table 1.

**Figure 7:**
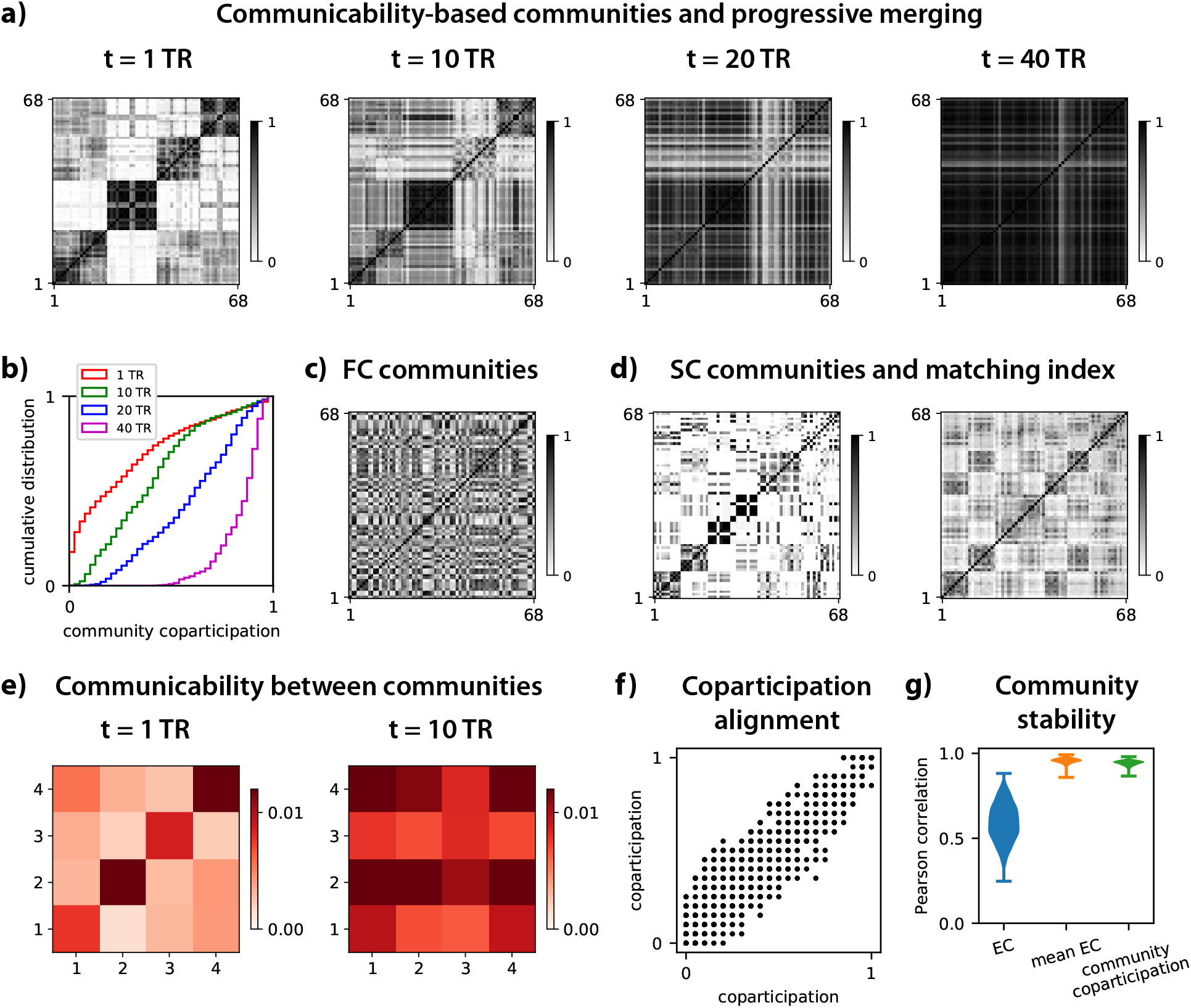
Communicability-based communities and homogenization of cortical activity. **a)** Each matrix represents the communities of cortical regions with strong communicability for four values of integration time *t*. Regions are paired when they have strong bidirectional communicability between them, see Eq. (7) for details. They correspond to averages over the 77 subjects and darker pixels indicate that the two regions (on the x- and y-axis) belong to the same community over all subjects. The region ordering is the same for all plots and is chosen to highlight the 4 communities (dark blocks on the diagonal). The x-axis indicate source ROIs and the y-axis target ROIs. **b)** Cumulative distribution of the community coparticipation values in panel a for the 4 matrices in panel a, illustrating the merging of communities. **c)** Similar plots to panel a with the FC-based communities. **d)** Similar plot (left panel) to panel a for SC-based communities obtained with the individual matrix with DWI values, see Methods. The right panel display the matrix of matching indices for SC, which measures the overlap between connected targets for each pair of ROIs. **e)** The matrices indicate the average communicability between the 4 communities obtained at *t* = 1 TR in panel a. **f)** Robustness of community detection. The plot represents the alignment of the communities for two subsets of 25% of the 77 subjects. Each plotted dot represents the coparticipation index that is a matrix element in Fig. 7a for *t* =1 TR. **g)** Pearson correlation coefficient between the MOU-EC of the 77 subjects, between the mean MOU-EC over each subject subset and of the community coparticipation values (100 repetitions using randomly 25% of the 77 subjects as in panel a, possibly with overlap).

We firstly compare the flow-based community matrices with the same algorithm applied on FC and SC, as classically performed. The resulting community matrices in Fig. 7c and d (left panel) are distinct from those obtained from communicability in several aspects. FC-based communities are less evident (with darker pixels on average), suggesting that even the strong overall correlations between ROIs do not adequately determine the functional communities. This comes for a large part because FC is a full matrix. We quantify the similarity between the community structures using the Pearson correlation on the vectorized coparticipation matrices in Fig. 7a-c-d. For communicability at *t* = 1 TR, this gives a Pearson correlation of 0.09 with the FC-based communities. In comparison we obtain 0.12 between SC- and FC-based communities. SC-based communities are more similar to the flow-based communities, with a Pearson 0f 0.54 between the coparticipation matrices. However, in each of the 4 groups, ROIs from distinct hemisphere are separated. This underlines the importance of the estimated MOU-EC weights, which give a different community structure compared to SC in spite of the same topology. The right matrix in Fig. 7c displays the pairwise matching indices calculated for the (binarized) SC. The matching index [69] quantifies the fraction of common neighbors two ROIs share and is thus often considered as a measure of “functional similarity”. Although the matching index is based on first-neighbor interactions, it leave the two hemispheres rather disconnectedwe find that this simplified estimate based on the topology alone satisfactorily predicts diffusion the propagation at early times described by communicability. Our results confirm that the correlation observed between input communicability and SC at the ROI level in Fig. 6 also applies to communities at the level of groups of ROIs. In contrast, the correlation between output communicability and FC does not yield a similar organization, with a similar low value to the comparison between SC and FC.

Another advantage of our approach is the quantitative description of the community merging over time, as illustrated by the distributions of coparticipation indices in Fig. 7d.

The communities are still differentiated at *t* ≤ 10 TR in Fig. 7a, but all of them except for the third one unite at *t* = 20 TR and they become a single community at *t* = 40 TR. The average communicability between the communities in Fig. 7e confirms this global pattern: It suggests that information is first integrated somehow independently within distinct communities, then is broadcasted to the whole network during the homogenization phase. These two modes of information integration are reminiscent of synchronization in networks for distinct values of the global coupling [70, 71], but it is important to note that they are supported by the same dynamical regime in our case; they simply occur at different timescales within the network response.

We verify the robustness of the community detection by performing the same community detection with two subsets of 25% of the 77 subjects. The left plot of Fig. 7f shows the correspondence between the coparticipation indices (matrix elements in Fig. 7f) for two subsets. From this we calculate the Pearson correlation coefficient to evaluate the alignment of the communities, which corresponds to 0.9 (green distribution in the right of Fig. 7g for 100 repetitions). This can be explained because the mean MOU-EC matrices over the subjects in each subset have a very similar structure (orange distribution), even though individual MOU-EC are more moderately aligned (blue distribution). Therefore, averaging over about 20 subjects reduces the subject variability, as well as noise related to the fMRI measurements.

### 3.5. Comparison of BOLD dynamics at different scales

Finally, we compare communicability between two parcellations to verify whether communicability corresponds to similar dynamics when changing scale. The Constellation parcellation is a refinement of the Desikan parcellation based on the anatomical fibers, dividing each of the original 68 ROIs into 3 to 5 new ROIs to obtain 237 ROIs with increased homogeneous SC within each ROI [52]. Although not identical, the reduced SC is thus very similar across the parcellations in Fig. 8a. The two FC matrices show perfect correspondence. Here we focus on an average model over subjects for MOU-EC, unlike the individual models used until now. The model can be satisfactorily fitted to the average FC0 and FC1 matrices (see the model error in Fig. 8b). As a sanity check, we verify that the average of individual MOU-EC matrices correlates with the MOU-EC of the average model with a Pearson of 0.56 (not shown), which is similar to the mean Pearson correlation between individual MOU-EC (blue distribution in Fig. 8b). The comparison between the average models of the two parcellations in Fig. 8c gives a Pearson correlation of 0.88, with p-value ≪ 10^−10^. The time constants *τ* are very close, 2.52 for Desikan and 2.53 for Constellation. With this we evaluate the total communicability as a proxy for the global network dynamics and find similar curves (Fig. 8d). These results indicate that consistent information can be obtained across scales, but we leave a deeper study for future work.

**Figure 8:**
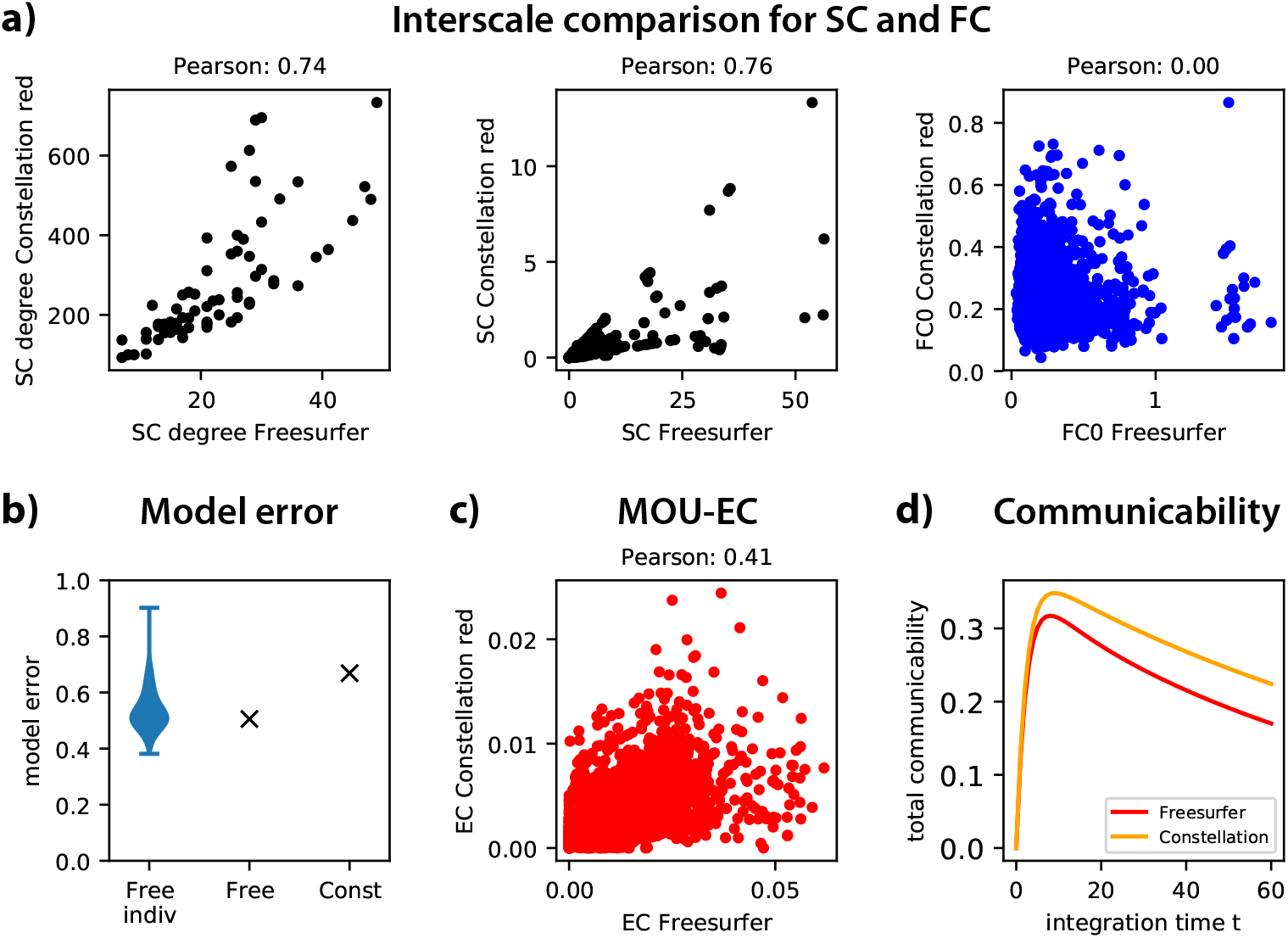
Comparison of communicability for two parcellations, Desikan and Constellation. In these plots the matrix values for Constellation with 237 ROIs are summed to reduce the matrix to the 68 ROIs of Desikan. **a)** Comparison of average SC degrees, SC values and FC values over subjects between the two parcellations. In the left plot, each black dot represents a ROI. In the middle and right plots each black/blue dot represents a link (pair of ROIs). **b)** Comparison of the model error for each subject (blue distribution) with the model error for the subject-average Desikan and Constellation models. **c)** Same as panel a for MOU-EC. d) Total communicability for the two parcellations.

## 4. Discussion

The present paper has introduced a network-oriented analysis of effective connectivity (MOU-EC) obtained from fMRI data, applying our recently-proposed formalism [37]. It demonstrates the flexibility of the framework that provides information at both local (single connections or ROIs) and global levels, allowing for a comprehensive description of the network going beyond a link-specific analysis. The pairwise interactions across time determined by the network dynamics are described by a family of matrices that incorporate network effects. Our results stress the importance of taking time into account to describe the brain communication in order to describe how “information” —here measured via the propagation of BOLD signals— is integrated in the network. Variability across subjects has not been explored here, but is of interest and will be studied in future work.

The results are summarized as plots on the cortical surface in Fig. 9. At the local scale, we have explored the functional roles of ROIs over time, e.g. broadcasters or listeners (Fig. 5a), which provides complementary information to FC and SC (Fig. 6). For instance, we have found that anatomical hub ROIs —precuneus, superior parietal and superior frontal cortex [7, 1]— are listeners that globally integrate information from other ROIs in the resting state, but without broadcasting much. During visual and memory tasks, they can become selective broadcasters to specific target ROIs [64]. Previous studies have shown that the precuneus also acts as a selective integrator during memory and cognitive flexibility tasks [65, 72]. It remains to be explored with diverse tasks how this gating of input/output information is implemented by high-levels ROIs, which can be related to the theory of the global workspace [28].

**Figure 9:**
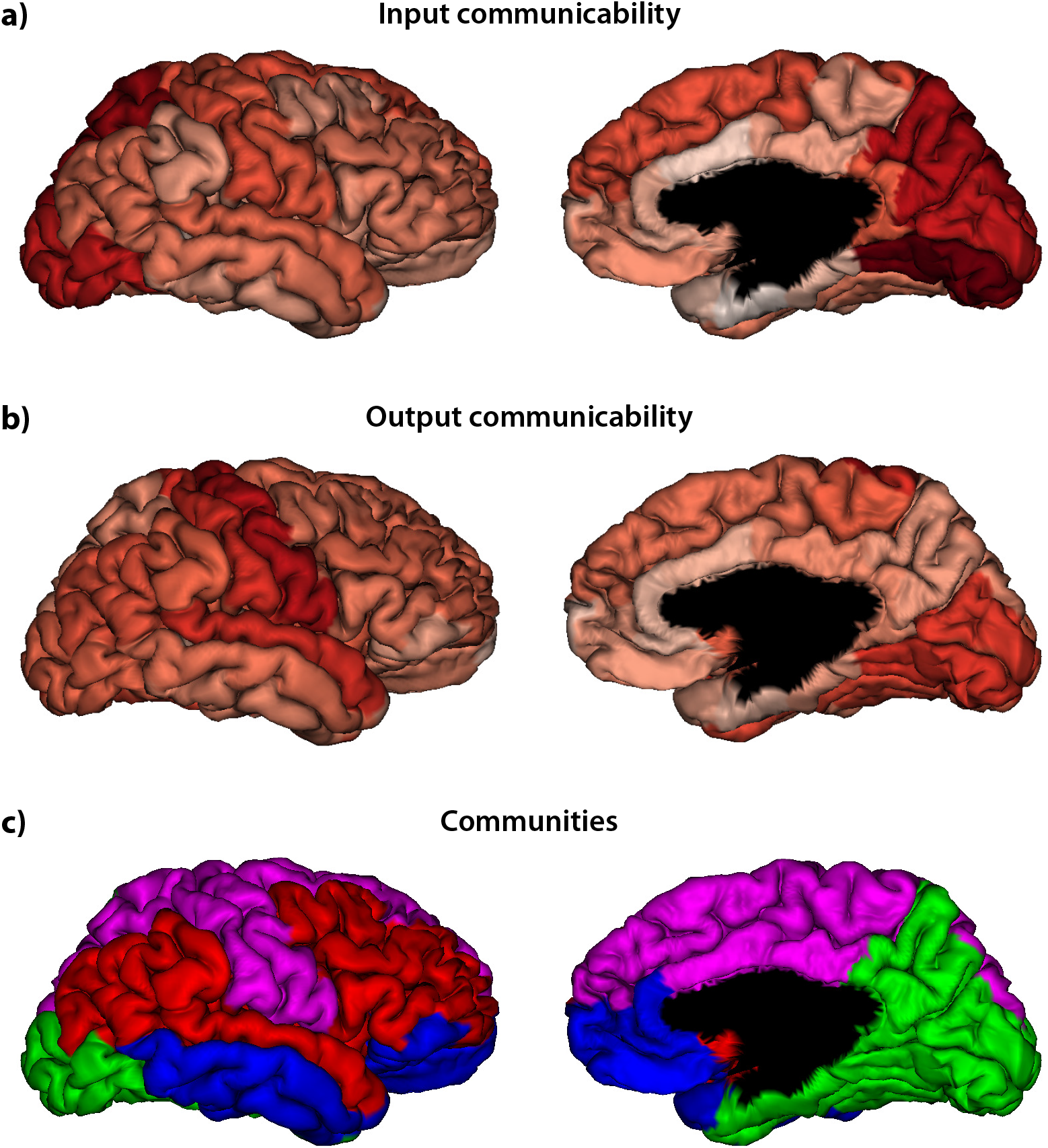
Summary of results. **a:** Mapping of the input communicability on the cortical surface (lateral and medial views of the right hemisphere). Darker red corresponds to higher value. **b:** Same as panel a for the output communicability. c: Mapping of the 4 communicability-based communities at *t* = 1 TR in Fig. 7a on the cortical surface. Each color represents a community.

At the global scale, the estimated BOLD dynamics exhibit a peak in the network response between 5 and 12 TR (Fig. 3c), while the maximum homogenization of dynamic communicability is achieved around 15 TR. This corresponds to community merging between 10 and 20 TR (Fig. 7a-b). These results speak to a multistage integration of information implemented by the cortical network —first locally within the communities, then globally. Here these two modes emerge as a natural consequence of the timescale separation due to information propagation within the network. In contrast, previous studies could only identify such modes by “artificially” setting the network into two very different dynamical states: either weakly or strongly synchronized [70, 71]. This flexibility may be interesting to quantify the notion of integration in networks that has attracted a lot of attention, in particular in neuroscience [29, 73, 28, 34, 32].

### 4.1. Model-based network descriptors for whole-brain dynamics

Our formalism opens a new dynamical perspective to interpret fMRI data, as compared to more “static” approaches using network theory that focused on FC or SC [7, 1]. The BOLD signals obtained from fMRI are “projected” on the model parameters. MOU-EC acts as a transition matrix, measuring the propagation of fMRI (fluctuating) activity across brain areas. In addition, the input properties are described via their (co)variance matrix Σ. In our model-based approach, the network analysis thus characterizes the brain *dynamics*, whose consistency from estimation to interpretation is a strength compared to previous studies in our opinion. As an example, FC-based analysis can be considered as a “projection” of BOLD signals on a graphical model, where the successive levels of BOLD activation (for all ROIs) are considered as i.i.d. variables over time. Likewise, sliding-window FC measures are typically considered as a pool of measures with no intrinsic temporal structure [22, 23, 24], which has also been used in other fields such as epidemic spreading [74]. Other alternatives to define states and transitions between them has used clustering techniques on the instantaneous phases obtained from the Hilbert transform [75] or HMMs to generate BOLD activation levels [26]. Although these approaches may be used to describe time-dependent interactions between ROIs, they do not capture the propagating nature of the BOLD signals, nor the network effect induced by these interactions. In addition, our approach conceptually differs from previous network studies that applied various types of nodal dynamics —such as Kuramoto oscillators and random walks— to static networks in order to reveal the properties of their complex topologies [38, 39, 40].

Importantly, the general concept of our dynamics-oriented network analysis is not tied to a specific model for whole-brain activity. The multivariate measure of communicability can be derived for other dynamic models provided their Green function is known or can be numerically evaluated, like the linearized dynamic causal model (DCM) used for resting-state fMRI data [41, 20, 76] or the multivariate autoregressive process [17]. This may lead to a revised formulation for Eq. (3). Nonetheless, a network analysis of the underlying connectivity —EC as discussed in a recent publication [77] or equivalent— only provides a snapshot view on the dynamics. Our results show that it corresponds to the initial integration time and that a more complete viewpoint is obtained when taking time into account (e.g. community merging). Other measures of directed connectivity can be used as matrix *A* to calculate communicability, such as Granger causality analysis [42, 43, 44]. However, this requires the choice of a leakage time constant *τ* in Eq. (1). In our case the knowledge about t comes for MOU-EC estimation procedure, which gives the full Jacobian *J*. Ensuring consistency from model estimation to interpretation is an important point in our opinion.

The presented framework allows for the analysis of connectivity at various scales. At the global level, the total communicability 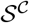 measures how the inputs circulate over the integration time via MOU-EC. The stabilization of 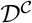 indicates the temporal horizon when the network interactions become most homogeneous. At the local scale, we have explored the functional roles of ROIs over time, e.g. broadcasters or listeners (Fig. 5a), which provides complementary information to FC and SC (Fig. 6). At an intermediate level Community detection uncovers the functional organization of ROIs (Fig. 7a). For the studied dataset, MOU-EC determines functional communities in a much more accurate manner than FC (Fig. 7c). This is an important point to analyze task-dependent communities, which is not possible using SC.

Another interesting aspect of our formalism is the quantitative comparison between network dynamics. With surrogate networks such as ring lattices or random networks, it reveals the properties of the estimated MOU-EC (Fig. 4). Here we have compared global measures: the total communicability 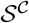 is the sum of all interactions at a given time and the diversity 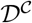 reflects their heterogeneity within the network. These time-dependent measures correspond to curves that are more or less distant, providing a metric to compare the data with specific topologies. Likewise, the comparison of the MOU-EC for two parcellations using communicability demonstrates a consistency across scales (Fig. 8).

In addition to the MOU-EC matrix *A* and the leakage related to *τ*, the MOU dynamics is determined by the properties of its fluctuating inputs (in purple in Fig. 3a), related to their (co)variance matrix Σ. The Σ estimated from data is task-dependent and off-diagonal elements may be tuned to model ROIs that experience cross-correlated noise [36]. To incorporate the effect of Σ in the network dynamics, another family of time-dependent matrices can be defined, which was name the flow [37]. In fact, dynamic communicability is a particular case of the flow for homogeneous input statistics (Σ_*ii*_ = 1 for all ROIs). Practically, the difference between the two measure is the following: Dynamic communicability describes how standard input fluctuations propagate throughout the estimated MOU-EC, whereas the flow quantifies the propagation of fluctuating activity for the brain dynamical “state” estimated over the whole session. Communicability is thus appropriate to study the propagation of perturbations in the network, whereas the flow is better suited when Σ depends on the subject’s condition as was shown for movie viewing versus rest [36].

### 4.2. Advantages and limitations of MOU-EC

The present application of dynamic communicability relies on the MOU process for the connectivity estimation. The MOU network corresponds to a non-conservative and stationary propagation of fluctuating activity, which was used to study network complexity [29, 30, 31, 32]. The MOU process has also been used in many other scientific disciplines [78, 79, 80], in particular for time-series analysis [81]. The viewpoint taken on the MOU process is a “noise”-diffusion network where fluctuating activity propagates, EC defining the input-output mapping [82]. Moreover, MOU-EC has been demonstrated to provide robust biomarkers for subject or task identification from BOLD signals [83].

When the MOU-EC estimation was developed, it aimed to solve the trade-off between robust estimation and application to large brain network (70+ ROIs) by using linear dynamics [18]. As mentioned earlier, the EC terminology was borrowed from the DCM literature because of the model-based aspect [15]. Several points were identified a few years ago when discussing the concepts behind effective connectivity and Granger causality analysis [57], in particular subsampling related to the low time resolution of BOLD signals and the hemodynamic response function. Because the MOU works in continuous time, it deals with the subsampling associated with the low temporal resolution of the BOLD signals [18]. This contrasts with estimation methods relying on the discrete-time multivariate autoregressive process that may be sensitive to subsampling for BOLD signals [43]. The absence of hemodynamics in MOU-EC is a crucial difference with DCM. An in-depth comparison between the two methods is left to future work and readers interested in whole-brain analysis involving hemodynamics are referred to recent extensions of the DCM [20, 76]. A related point concerns the single time constant *τ* for all ROIs chosen for each subject, which was motivated from the analyzed data. Further study —possibly with more datasets— is necessary to explore the possibility of finessing the model with ROI-specific *τ*. Recall also that any interpretation of BOLD in term of brain communication relies on the assumption that changes in neural activity are reliably reflected in the BOLD signals, which is under debate [84, 85, 86].

The model optimization enforces a diagonal Σ, which has consequences on the estimated MOU-EC. This observed in a previous study for task-evoked activity [36], but not consistently explored so far. In the present version the MOU-EC does not incorporate common inputs, which means that the *A* estimates may compensate with stronger weights to explain the observed correlations in FC. However, there is no whole-brain dynamic model that has quantitatively addressed this difficult issue in depth so far in our knowledge.

## Acknowledgements

This work has been supported by the European Union’s Horizon 2020 research and innovation programme under Grant Agreement No. 720270 (HBP SGA1) and No. 785907 (HBP SGA2). MG also acknowledges funding from the Marie Skłodowska-Curie Action (Grant H2020-MSCA-656547) of the European Commission. GZL, NEK and GD acknowledge funding from the European Union’s Horizon 2020 research and innovation programme under Grant Agreement No. 720270 (HBP SGA1). GD also acknowledges funding from the Spanish Research Project (No. PSI2013-42091-P). NEK acknowledges support from the “MOVE-IN Louvain” fellowship co-funded by the Marie Skłodowska-Curie Action of the European Commission.

The “CONNECT/Archi Database” is the property of the CEA I2BM NeuroSpin centre and was designed under the supervision of Dr Cyril Poupon and Dr Jean-François Mangin. It was funded by the Federative Research Institute 49, by the HIPPIP European grant, and the European CONNECT project (http://www.brain-connect.eu). Acquisitions were performed by the scientists involved in the Multi-scale Brain Architecture research program of NeuroSpin and by the staff of the UNIACT Laboratory of NeuroSpin (headed by Dr Lucie Hertz-Pannier), under the ethical approval CPP100002/CPP100022 (principal investigator Dr Denis Le Bihan). Access to the database can be requested from.

